# The apical Ciliary Adhesion complex is established at the basal foot of motile cilia and depends on the microtubule network

**DOI:** 10.1101/2022.06.28.497743

**Authors:** Maria Chatzifrangkeskou, Paris A. Skourides

## Abstract

The Ciliary Adhesion (CA) complex forms in close association with the basal bodies of cilia during the early stages of ciliogenesis and is responsible for mediating complex interactions with the actin networks of multiciliated cells (MCCs). However, its precise localization with respect to basal body accessory structures and the interactions that lead to its establishment in MCCs are not well understood. Here, we studied the distribution of the CA proteins using super-resolution imaging and possible interactions with the microtubule network. The results of this study reveal that the apical CA complex forms at the distal end of the basal foot and depends on microtubules. Our data also raise the possibility that CAs may have additional roles in the regulation of the organization of the microtubule network of MCCs.

## Introduction

Cilia are microtubule-based protrusions, and they are classified into primary and motile cilia that differ in structure and function^1,2^. Most cells develop a single, non-motile, primary cilium for sensing and transducing mechanical and chemical signals. Motile cilia, on the other hand, beat in a coordinated and polarized manner to generate fluid flow. For instance, multiciliated cells (MCCs) contain hundreds of motile cilia that drive cerebrospinal fluid flow in the brain and oocyte transportation along the oviduct and they are required in the respiratory tract for the clearance of mucus that traps inhaled particles and pathogens. Therefore, defects of the cilia-generated fluid flow result in increased risks of hydrocephalus, and female infertility, chronic recurrent respiratory infections, and lead to diseases such as primary ciliary dyskinesia (PCD)^3–5^.

Cilia are assembled from basal bodies (BBs), which are ninefold symmetrical microtubule (MT)-based structures. In MCCs, BBs are polarized with the basal feet projecting towards the direction of the ciliary flow. Loss of the basal foot leads to ciliary disorientation^6^. The basal foot functions as an MT-organizing center and its proteins are organized in three structural regions: region I connects the basal foot to the microtubule triplets of the basal body and is likely providing mechanical support during effective stroke^6,7^; region II or scaffolding region consists of proteins that serve in the assembly of the basal foot, such as ODF2^6,8,9^, Ninein^10^, galactin-3^7^, CEP19^11^, CEP128^12^, centriolin^13^ and CC2D2A^14^; region III or microtubule-anchoring region corresponds to the basal foot cap which is comprised of a protein complex that anchor or nucleate microtubules. Specifically, in motile cilia, the cap contains among other proteins NEDD1 and γ-tubulin that develop the γTuRC complex, necessary for microtubule nucleation^15^. CEP170 is a basal foot cap protein required to connect basal bodies to the microtubule network^16,17^. In addition, both z- and ε-tubulin localize at the basal foot cap in MCCs^18^. Interestingly, depletion of z-tubulin does not disrupt the connection of MT network to the basal foot, but it perturbs the assembly of the apical and subapical actin networks, suggesting that the basal foot mediates basal body interactions with the actin cytoskeleton^18^.

The embryonic epidermis of *Xenopus laevis* has an array of MCCs providing an ideal model to study cilia development and function. Our previous work using *Xenopus laevis* led to the discovery of a novel ciliary structure, termed ciliary adhesions (CAs)^19^. CAs are multiprotein complexes which associate with the basal bodies of cilia and are involved in the interactions between basal bodies and the actin cytoskeleton during MCC differentiation, as well as in mature MCCs. Four protein members of the CA complex were identified: FAK, Paxillin, Vinculin and Talin; all being well-characterized members of the Focal Adhesion (FA) multiprotein complex. CAs form in two distinct regions sub-apically at the end of the striated rootlet and apically adjacent to the basal bodies, however their precise positioning in relation to basal body accessory structures remains unexplored. In the present study, we performed a more detailed localization analysis of CA proteins using super-resolution microscopy to increase imaging resolution and improve localization accuracy. CA sub-ciliary positioning was determined in reference to proximal and distal markers of the basal foot. We, thus, identified CA members as basal foot proteins aiding future investigations on the role of basal foot components in the biology of cilia.

## Results

### CAs associate with microtubules

Based on our previous observations, the CA signal is concentrated adjacent to the basal bodies and opposite to the rootlets, approximately at the area where the basal foot forms and at a site where electron microscopy analysis identified interactions between basal bodies and the actin cytoskeleton^20^. Since TEM analysis confirmed that the basal foot directly interacts with cortical microtubules in mature MCCs^17^, we initially analyzed the localization of the CA protein, FAK, in relation to the apical microtubule network in *Xenopus* epidermal MCCs. Whole-mount immunostaining of Xenopus embryos with an antibody against β-Tubulin gave strong staining of cilia, compromising our ability to image the intracellular MT network in mature MCCs. To overcome this, a deciliation strategy was employed to eliminate cilia-associated signals and facilitate visualization of intracellular MTs. Embryos injected with GFP-FAK/RFP-Centrin were deciliated using calcium chloride (CaCl_2_), a known deciliating agent, for 1 min, and fixed with ice cold methanol which was the only fixative tested that preserved apical CA localization and retained a pattern similar to that obtained via live imaging. Aldehyde based fixation of CA components generates artifacts with the majority of the signal from CA proteins concentrated on the striated rootlet (Fig S1). As shown in Fig 1A, FAK signal is overlapping with β-Tubulin, indicating partial colocalization of the CA complex with the microtubule network. To explore the possibility that the establishment of the CA complex may depend on interactions with the apical microtubule network, GFP-FAK/RFP-Centrin embryos were treated with 1 μM nocodazole from stage 23-29, which has been previously shown to disrupt the highly organized apical microtubule network in MCCs^21^. Interestingly, nocodazole treatment markedly affects the CA localization of FAK and its association with the basal bodies (Fig 1B). This suggests that FAK recruitment at CAs is at least partially dependent on the apical microtubule network. In addition the position of the FAK signal with respect to the BB is affected in nocodazole treated embryos (Fig 1B). This may indicate structural changes of the BF on which CAs form or repositioning of the CA on said structure.

**Figure 1:**
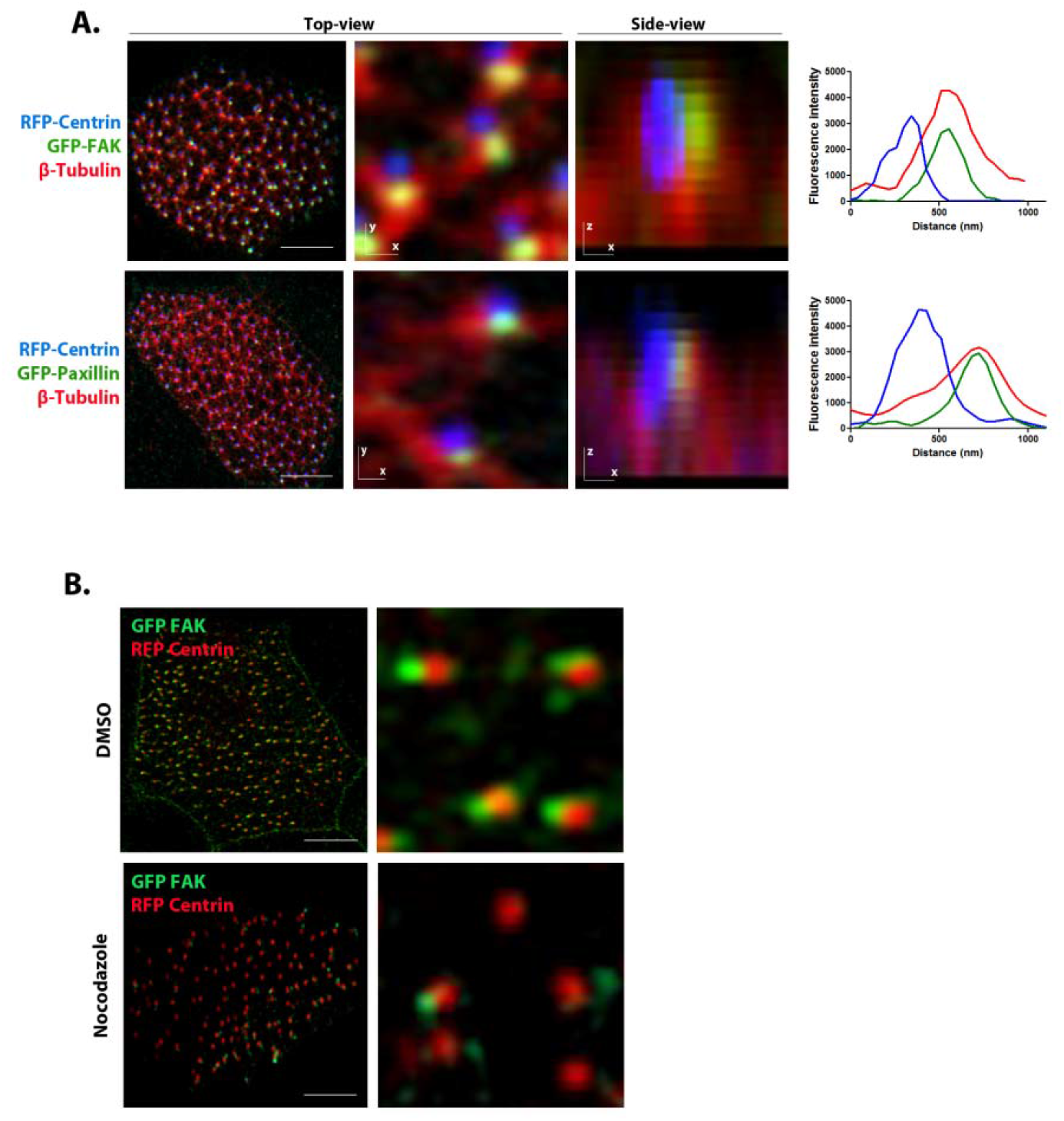
CAs associate with microtubules. A. Co-expression of RFP-Centrin and GFP-FAK followed by β-tubulin staining in deciliated MCCs. Scale bars, 5l.1µm. B. Embryos injected with RFP-Centrin and GFP-FAK were treated with 0.1% DMSO or 1μM Nocodazole from stage 23-29.

### Mapping of basal foot proteins in *Xenopus* motile cilia

Given the proximity of CAs to BFs and BBs, we decided to map the precise position of CAs within the BF structure. However, the molecular architecture of the basal foot in the motile cilia of *Xenopus* is still lacking so we first used a combination of antibodies and fusions of proteins that have been assigned to the proximal (GFP-CEP128, GFP-ODF2, CEP19, GFP-Gal3) and distal region (GFP-CEP170, mCherry z-Tubulin, mOrange γ-Tubulin) of the basal foot in mammalian to generate a map of the *Xenopus* basal foot using super-resolution microscopy. We imaged most of the reported basal foot proteins in combination with mTagBFP2 Centrin that serves as a marker for basal bodies and quantitatively mapped protein positions relative to the center of the basal body (Fig 2). The mRNA of basal foot markers together with Centrin mRNA was introduced via targeted microinjections at the ventral blastomeres of 4-cell stage embryos. The basal foot region-II components CEP128, ODF2 and CEP19 showed similar distances from the basal body center. Similarly to human primary cilia, Gal3 resides in the bridging region between regions II and III, while zeta- and γ-Tubulin are the most distant from the basal body center, consistently with their association with MTs. The distribution of CEP170 in close proximity to the region III of basal foot is consistent with its ability to bind MTs^22^. Overall, our data showed that the distribution of basal foot proteins in *Xenopus* MCCs is consistent with the previous reported positions in human motile cilia^15^.

**Figure 2:**
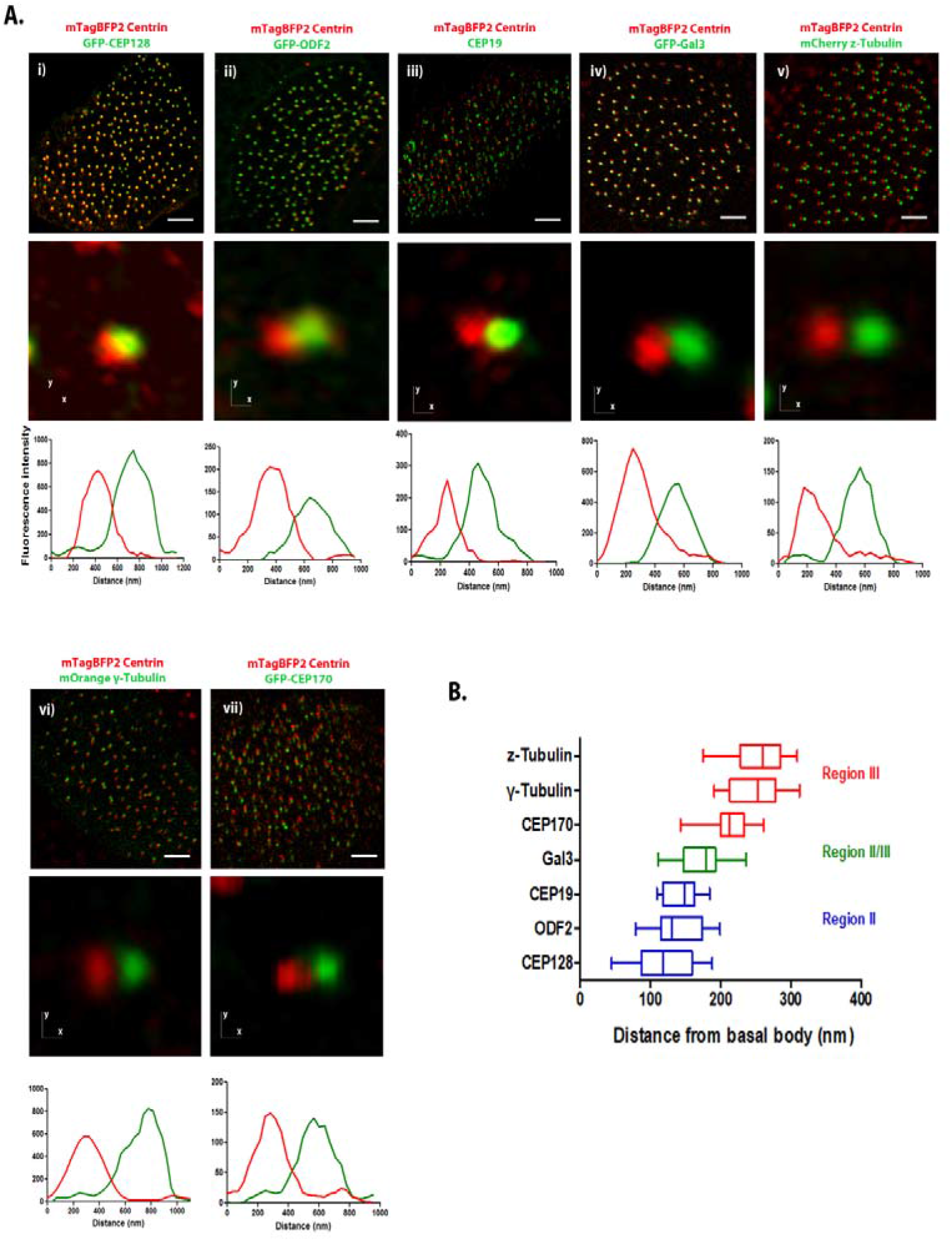
Mapping of basal foot proteins in *Xenopus* motile cilia. **A**. i) Optical section of a multiciliated cell expressing mTagBFP2-Centrin and GFP-CEP128. ii) Optical section of a multiciliated cell expressing mTagBFP2-Centrin and GFP-ODF2. iii) Optical section of a multiciliated cell expressing mTagBFP2-Centrin and stained with an antibody against CEP19. iv) Optical section of a multiciliated cell expressing mTagBFP2-Centrin and GFP-Gal3. v) Optical section of a multiciliated cell expressing mTagBFP2-Centrin and mCherry z-Tubulin. vi) Optical section of a multiciliated cell expressing mTagBFP2-Centrin and mOrange γ-Tubulin. vii) Optical section of a multiciliated cell expressing mTagBFP2-Centrin and GFP-CEP170. Scale bars represent 5 μm. B. Box plot of distance measurements of basal foot proteins in epidermal MCCs in *Xenopus*.

### FAK is localized at the distal end of the basal foot

Using this super-resolution map as a guide, we next examined CA localization in relation to proteins that have been assigned to the proximal region of the basal foot. Analysis of the relative distribution of the GFP-ODF2 and GFP-128 in live imaging of individual basal bodies of MCCs revealed that none of these region II basal foot markers co-localized with mKate2-FAK (Fig 3A). Lateral views and intensity profiles of the magnified inset show distinct peaks of the fluorescent signals.

**Figure 3:**
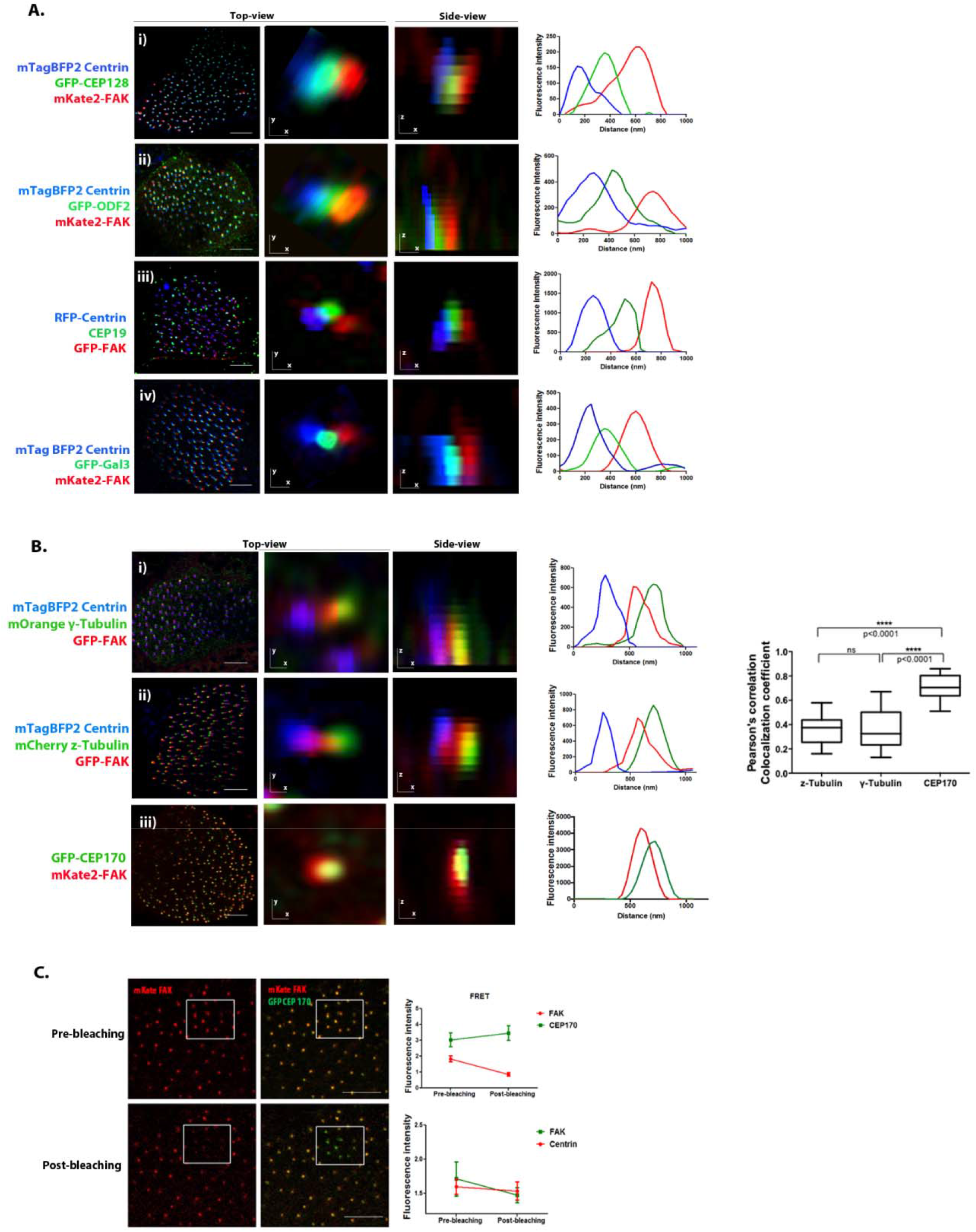
FAK is localized at the distal end of the basal foot. A. i) Confocal optical section of a multiciliated cell expressing GFP-CEP128, mKate2-FAK and mTagBFP2-Centrin. High-magnification view of a representative basal body and orthogonal (x-z) image stacks of a multiciliated cell are shown. ii) Confocal optical section of a multiciliated cell expressing GFP-ODF2, mKate2-FAK and mTagBFP2-Centrin. Right: High-magnification view of a representative basal body and orthogonal (x-z) image stacks of a multiciliated cell are shown. iii) Immunofluorescence staining of endogenous CEP19 in RFP-Centrin/GFP-FAK expressing multiciliated cells. iv) Optical section of a multiciliated cell expressing GFP-Gal3, mKate2-FAK and mTagBFP2-Centrin. Top view and orthogonal projection (x-z) are shown. Scale bars represent 5 μm. B. i) Representative optical section of a multiciliated cell expressing GFP-FAK, mOrange γ-Tubulin and mTagBFP2-Centrin. Top view and orthogonal projection (x-z) are shown. ii) Optical section of a multiciliated cell expressing GFP-FAK, mCherry z-Tubulin and mTagBFP2-Centrin. Top view and orthogonal projection (x-z) are shown. iii) Representative optical section of a multiciliated cell expressing GFP-CEP170 and mKate2-FAK. Higher top-view magnification of a single basal body and a 3D reconstruction (x-z) of optical sections is shown. Scale bars represent 5 μm. Pearson’s correlation coefficient graph representing colocalization of FAK with z-Tubulin, γ-Tubulin and CEP170. Higher values indicate stronger colocalization. C. Multiciliated cell expressing GFP-CEP170 and mKate2-FAK before and after acceptor photobleaching. Quantification of the acceptor (mKate2) and donor (GFP) fluorescence intensity following photobleaching. Control experiments with GFP-Centrin as the donor and mKate2-FAK as an acceptor were carried out. Scale bars represent 10 μm.

In addition, distribution analysis of FAK with respect to centrin and CEP19 suggested a more distal localization of the CAs (Fig 3A). We, therefore, next studied the localization of FAK in relation to Galectin-3, which has been shown to have a broad distribution, in primary cilia, extending toward the bridging region between regions II and III of the basal foot^15^. However, fluorescence intensity profiles showed that FAK is located more distally to both bridging region markers. Taken together these data suggest that FAK is located more distally than the known markers of the proximal regions of the basal foot, suggesting that the CA complex may form at the distal region III.

We next investigated whether the CA complex is established at the distal end of the basal foot in multiciliated cells of the tadpole epidermis. We analyzed the distribution of FAK relative to the microtubule nucleator γ-Tubulin, known to localize to the basal foot cap in MCCs^7,23^. Expression of GFP-FAK in MCCs was found to partially overlap with γ-Tubulin, in line with our previous data suggesting that CA member proteins localize in the vicinity of the basal foot cap (Fig. 3B). Next, we analyzed the distribution of FAK relative to Centrin and z-Tubulin, another component of the basal foot cap^18^. Similarly, fluorescence intensity profiles reveal partial co-localization of z-tubulin and FAK, confirming that CAs are formed in close proximity to the basal foot cap.

To further investigate this, we co-expressed mKate2-FAK and GFP-CEP170, a protein located in region III^16^. As shown, the signal from FAK overlaps with the signal from CEP170 (Fig. 3B). In fact the two signals display a higher colocalization coefficient compared to any other marker tested. The above data show that CAs form at region III of the basal foot and the clear colocalization between FAK and CEP170 raises the possibility that the two interact. In order to examine this possibility, we co-expressed GFP-CEP170 (donor) with mKate2-FAK (acceptor) and carried out acceptor photobleaching experiments to examine the possibility of intermolecular fluorescence resonance energy transfer (FRET). These experiments show that FRET is taking place between GFP-CEP170 and mKate2-FAK in MCCs (Fig 3C). Control experiments using centrin GFP as the donor and mKate2-FAK as an acceptor fail to detect FRET, as expected, and show that FAK does not interact with centrin. Thus, we conclude that the CA complex is established at the distal end of the basal foot in MCCs and may depend on an interaction between FAK and CEP170.

### FAK depletion induces defects of the MCC microtubule network

The localization data raised the possibility that CAs could participate in the formation of the basal foot in *Xenopus* MCCs. To address this possibility, we injected *Xenopus* embryos with a previously characterized translation-blocking morpholino (MO) to knock down FAK expression in multiciliated cells^19,24,25^. We examined the localization of basal foot markers by co-expressing fluorescently tagged z-Tubulin and Galectin-3, after basal body orientation is established (stages 30-35). As shown the basal foot is established in FAK morphants, as the basal foot cap component z-Tubulin and the proximal basal foot protein Galectin-3 were still present (Fig 4A). Interestingly, the distance between the proximal and the distal basal foot markers was increased in FAK morphants compared to controls suggesting that the basal foot structure may be affected leading to longer structures.

**Figure 4:**
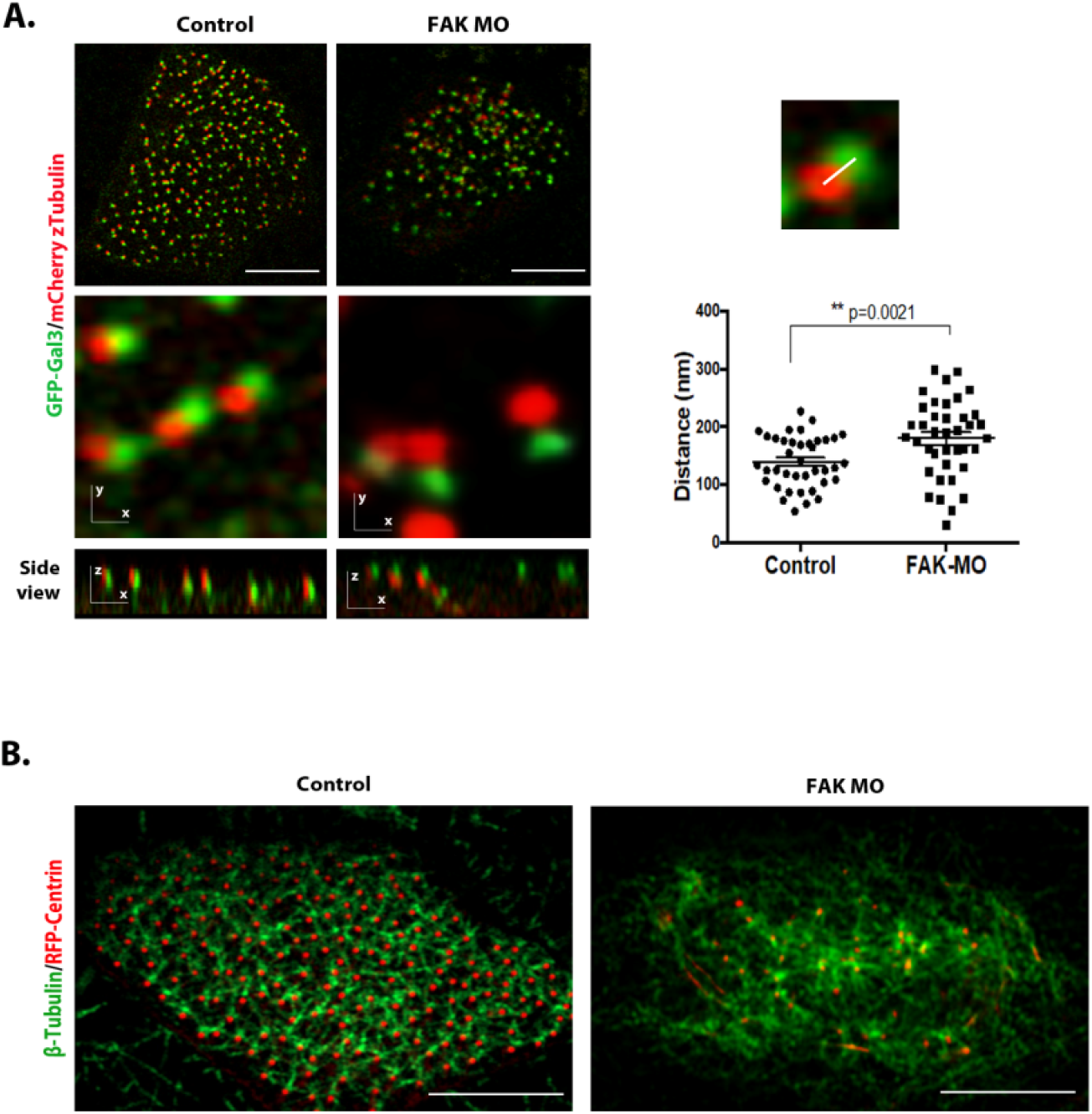
FAK depletion induces defects of the MCC microtubule network. A. *Left:* GFP-Galectin 3 and mCherry z-Tubulin-expressing control and FAK morphant multiciliated cells. scale bars, 5⍰µm. *Right:* Quantification of the distance between GFP-Galectin 3 and z-Tubulin. Error bars represent mean ± SEM. B. Microtubule network in control and FAK morpholino-injected multiciliated cells was stained with an antibody against β-tubulin. Embryos were deciliated and imaged at stage 35.

The partial co-localization of the CAs with the basal foot cap, which acts as a microtubule-organizing center (MTOC) that participates in microtubule nucleation and/or stabilization raises the possibility that CAs might be involved in these processes. Therefore, we also analyzed the apical microtubule network in FAK morphants. Depletion of FAK in MCCs leads to severe disruption of the apical microtubule network (Fig 4B). However we cannot exclude the possibility that this is a secondary effect of the basal body docking defects observed in FAK morphants.

Overall the above data suggest that FAK downregulation impacts the structure of the basal foot given the changes in the spatial relationship of Gal3 and z-Tubulin and elicits severe defects of the apical microtubule network which however could be secondary to the failure of basal bodies to dock.

### Hierarchical association of CEP170 and FAK with basal bodies

The CA members FAK and paxillin showed significant but not perfect colocalization (Fig S2). The close spatial relationship between CAs and CEP170 as well as the possible interaction between CA members and this protein raises the possibility that CEP170 may be mediating interactions between BBs and CA proteins. Since FAK is present during basal body migration^19^, we examined the temporal relationship between FAK, Paxillin and CEP170 with respect to their association with the BBs. As shown in Fig 5A, CEP170 is associated with the basal bodies, marked by mTag-BFP2 centrin, from stage 16, during ciliated cell intercalation. At this stage when radial intercalation of MCCs begins, although CEP170 was clearly associated with the basal bodies, FAK was only weakly detected (Fig 5B). At stage 18, when centrioles start migrating towards the apical surface, FAK colocalizes with CEP170 and is closely associated with the basal bodies. At stage 20 when cells begin to undergo ciliogenesis, FAK was visible near mature basal bodies docked at the apical surface. We went on to compare the temporal association kinetics of FAK to those of paxillin. Pearson’s correlation coefficient analysis shows that unlike FAK, paxillin is clearly associated with the BBs at the earliest time points at stage 16 similar to CEP170 (Fig 5D). This suggests that paxillin may be recruited at the BBs via CEP170 and subsequently recruits FAK to form the CA complex in agreement with previous data showing that FAK relies on its interaction with paxillin for CA localization. Future work will focus on the role of CEP170 on the establishment of the apical CA complex.

**Figure 5:**
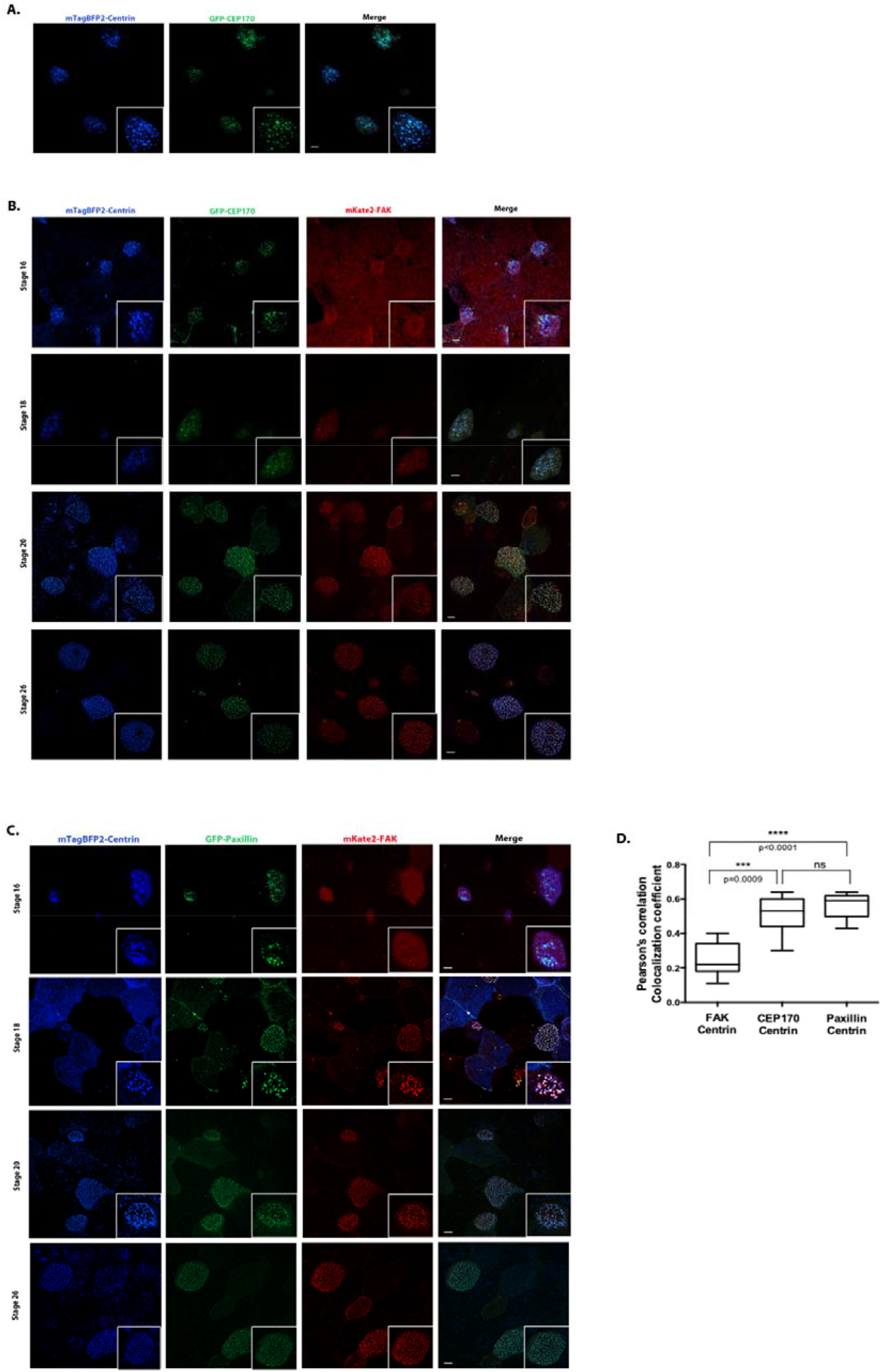
Hierarchical association of CEP170 and FAK with basal bodies. A. Intercalating ciliated cells of stage 17 *Xenopus* embryos coexpressing mTag-BFP2-Centrin and GFP-CEP170. Scale bars represent 5 μm. B. Intercalating ciliated cells of *Xenopus* embryos coexpressing mTag-BFP2-Centrin, mKate2-FAK and GFP-CEP170. Embryos were imaged at the indicated stages. Scale bars represent 5 μm. C. Intercalating ciliated cells of *Xenopus* embryos coexpressing mTag-BFP2-Centrin, mKate2-FAK and GFP-Paxillin. Embryos were imaged at the indicated stages. Scale bars represent 5 μm. D. Pearson’s correlation coefficient graphs representing the colocalization of mKate2-FAK, GFP-CEP170 and GFP-Paxillin with mTagBFP2-Centrin.

## Discussion

We previously identified a novel role of FAK, as a member of CA complexes in Xenopus epidermal MCCs^19^. Loss of FAK in these cells leads to impaired ciliogenesis due to defective association of the basal bodies with the actin cytoskeleton. In MCCs, CAs are located posterior to the basal bodies, with respect to the tadpole’s A-P axis, where the basal foot resides, but its precise localization was not known. Taking advantage of super-resolution microscopy, we were able to compare the localization of FAK to that of BF markers and define the region at which apical CAs form with high spatial resolution. We found that FAK is localized close to the basal foot cap (Fig 6). More specifically, we found that FAK and the region III-associated protein CEP170, spatially overlap and display a very high colocalization coefficient. This shows that apical CAs form at region III of the basal foot.

**Figure 6:**
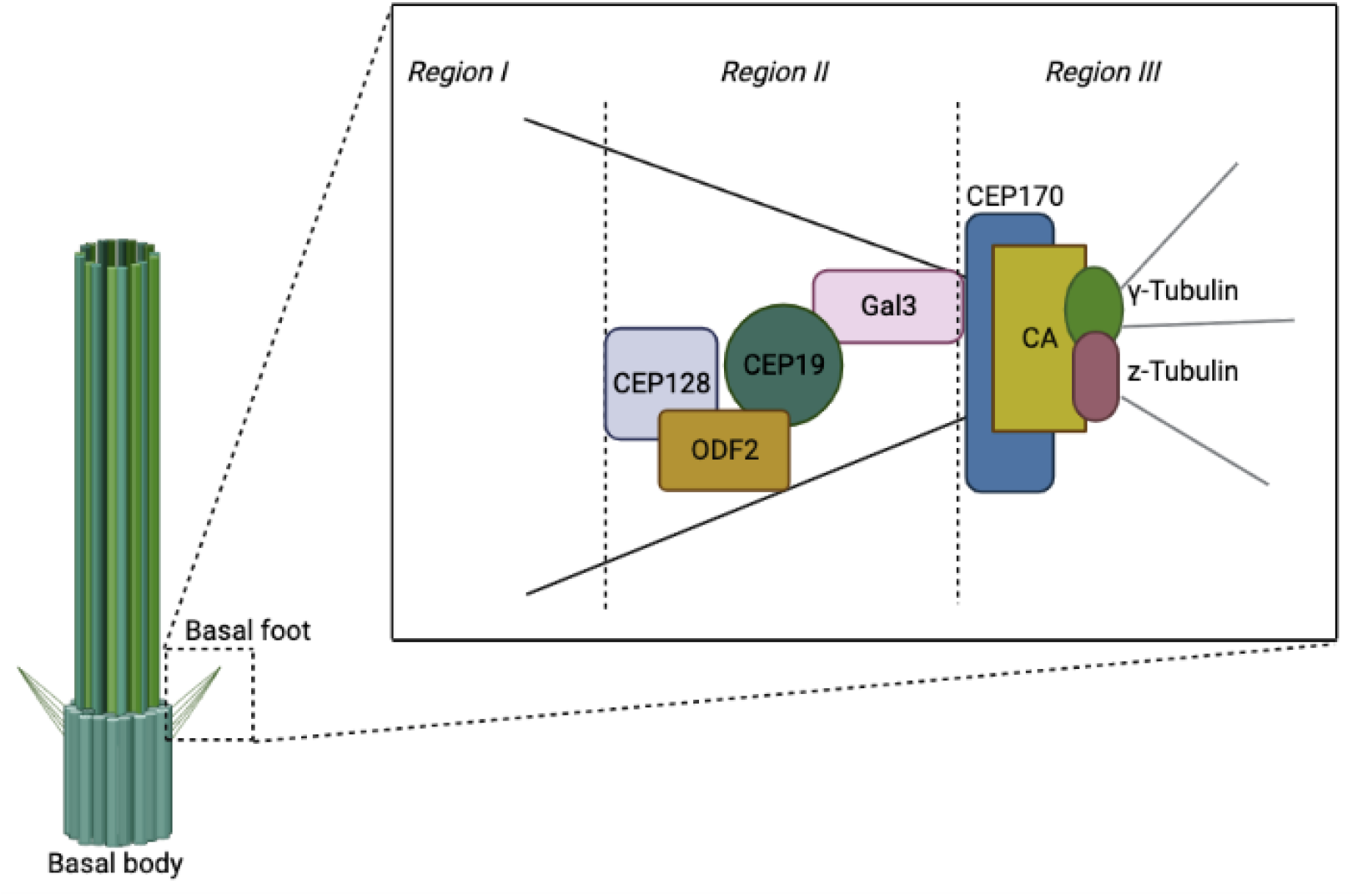
CA localization at the distal region of the basal foot. Cartoon depiction of CA localization at the basal foot. Basal foot components are organized into spatially distinct regions with different functions (region I, basal body anchoring; region II, scaffolding; region III, MT anchoring).

The localization of FAK in the CA complex strongly depends on the FAT domain and its interaction with Paxillin, similarly to FAK’s localization at FAs. In fact, the FAK I936E/I998E mutant that does not bind paxillin fails to be recruited at CAs and fails to rescue FAK morphants showing that the interaction with paxillin is required for both localization and function^26^. In line with these results, we observed that Paxillin appears to be associated with basal bodies in multiciliated cells earlier than FAK. Interestingly, CEP170 is also associated with the BBs at early stages similar to paxillin raising the possibility that paxillin is recruited at the basal foot by CEP170 and subsequently recruits FAK to form the CA complex.

Loss of the basal foot in mice leads to respiratory manifestations indicative of PCD. In tracheal MCCs, loss of the basal foot results in disruption of the microtubule apical network and disorientation of basal bodies^6^. The localization of FAK at the basal foot cap suggests that CAs might be involved in the establishment or maintenance of apical microtubule network. FAK morphants display longer basal feet and altered microtubule network (Fig. 3), suggesting that CAs may play a role in regulating the BF structure. Despite the critical role of the basal foot in multiciliated cell function, its molecular organization in *Xenopus* remains to be elucidated. In addition to defining the precise localization and dissecting the role of the CA protein FAK, we also provide a molecular map of known basal foot components (ODF2, CEP128, Galectin 3, CEP19, Ninein, CEP170, z-Tubulin and γ-Tubulin) in *Xenopus* motile cilia. Using super-resolution imaging, we showed that the basal foot in *Xenopu*s MCCs bears a high level of similarity with the basal foot of human airway multiciliated cells^15^. This high level of conservation between the human and the *Xenopus* basal foot reaffirms *Xenopus* as a valuable model system for the study of motile cilia.

## Methods

### Embryo Manipulations, Microinjections, and Lysates

Female adult *Xenopus laevis* were ovulated by injection of human chorionic gonadotropin. Eggs were fertilized *in vitro*, de-jellied in 1.8 % cysteine (pH 7.8), and embryos were maintained in 0.1× Marc’s modified ringers (MMR) and staged according to Nieuwkoop and Faber. Microinjections were performed in 4% Ficoll in 0.33× MMR at the ventral blastomeres of 4- or 8-cell-stage embryos to target the epidermis. For MO experiments, we injected 12 ng of the FAK morpholino per blastomere (at 8-cell stage embryos). For live imaging, embryos were anesthetized in 0.01% benzocaine in 0.1× MMR and immobilized in silicone grease wells on glass slides.

### DNA Constructs and Morpholino Oligonucleotide

All plasmids were transcribed into mRNA using the SP6 or T7 mMessage mMachine kit® (Invitrogen) and used for microinjections. The GFP-CEP128 plasmid was kindly provided by Lotte Bang Pedersen’s laboratory. The sequence of FAK MO used in our experiments is TTGGGTCCAGGTAAGCCGCAGCCA^19,26^.

### Immunostaining

Embryos were allowed to develop to the appropriate stage and then imaged live or fixed in Methanol at -20°C for 2 hours and gradually rehydrated with 75% methanol-25% MEMFA (10X: 1M MOPS, 20mM EGTA, 10mM MgSO4, 38% Formaldehyde), 50% methanol-50% MEMFA, 25% methanol-75% MEMFA and 100% MEMFA for 10 minutes each. Embryos were then permeabilized in PBDT (1× PBS + 0.5% Triton X-100 + 1% DMSO) for several hours at room temperature and blocked in PBDT + 10% donkey serum for 1 h at room temperature. Primary antibodies were then added (in block solution). Primary antibodies used were: β-tubulin (Hybridoma), γ-Tubulin (Abcam) and CEP19 (Proteintech). The embryos were incubated in the antibody solution overnight at 4°C. The next day embryos were washed in PBDT and then incubated in secondary antibodies diluted in the blocking solution at room temperature for 1 h. Imaging was performed on a Zeiss LSM 900 laser scanning confocal microscope with Airyscan imaging system. Image processing and colocalization analysis was done using the Zeiss ZEN 2.3 software.

### Acceptor Photobleaching FRET

FRET experiments were accomplished using a laser scanning confocal microscope (Zeiss LSM 710) with a Plan-Apochromat 63×/1.40 Oil DIC M27 objective lens (Zeiss). A 543 nm laser was used for acceptor (mKate-FAK) photobleaching within a region of interest (ROI). One frame was acquired as a pre-bleaching control, and the ROI was bleached within one frame. Zeiss Zen 2010 software was used for FRET analysis.

## Author Contributions

M.C carried out the experiments and data analysis. M.C and P.A.S wrote the manuscript.

## Acknowledgements

We thank Dr. Pedersen for kindly providing the GFP-CEP128 plasmid.

