## Supplementary figures and images for "The apical Ciliary Adhesion complex is established at the basal foot of motile cilia and depends on the microtubule network"

### (Fig S1).

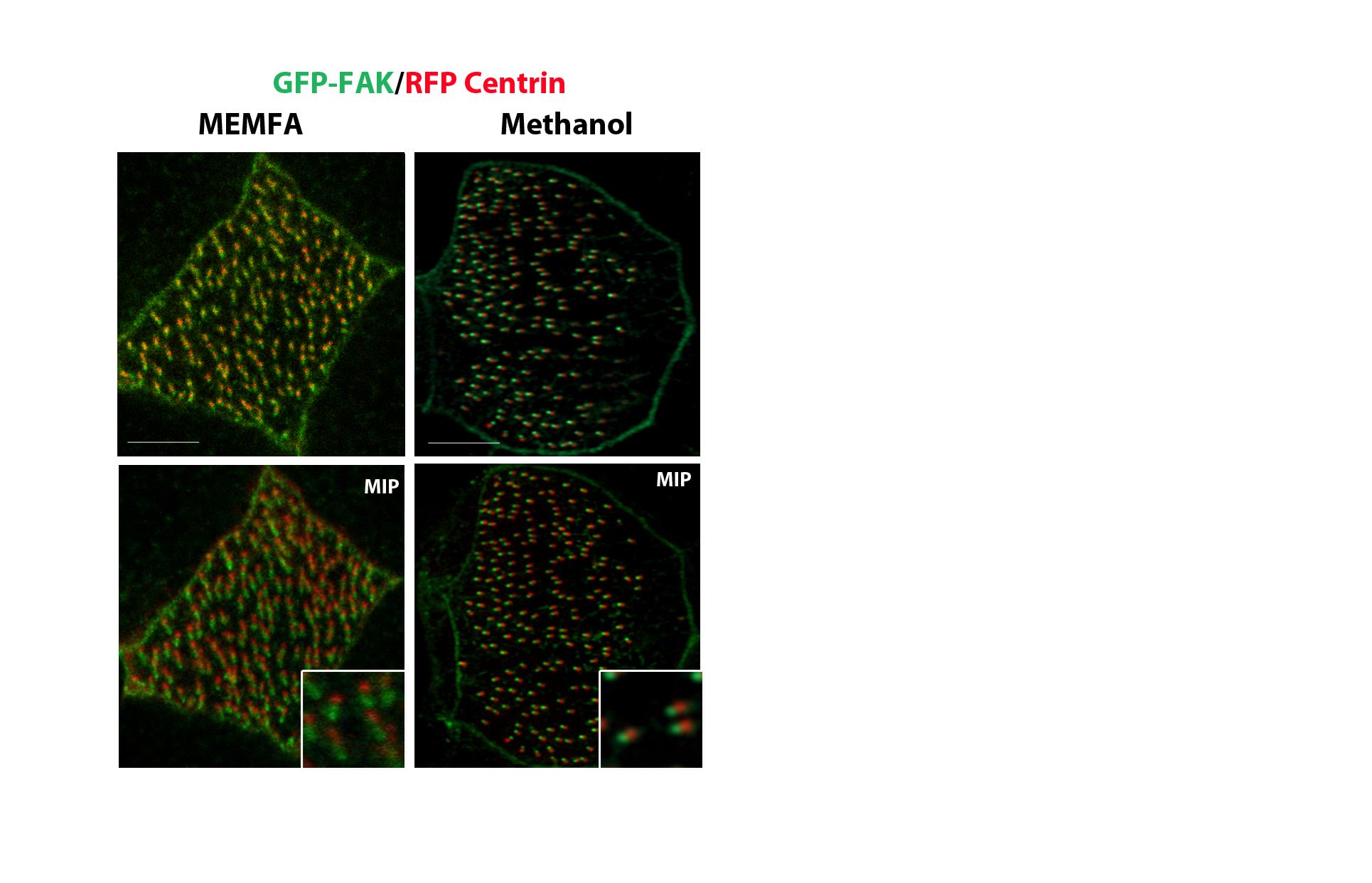

### (Fig S2).

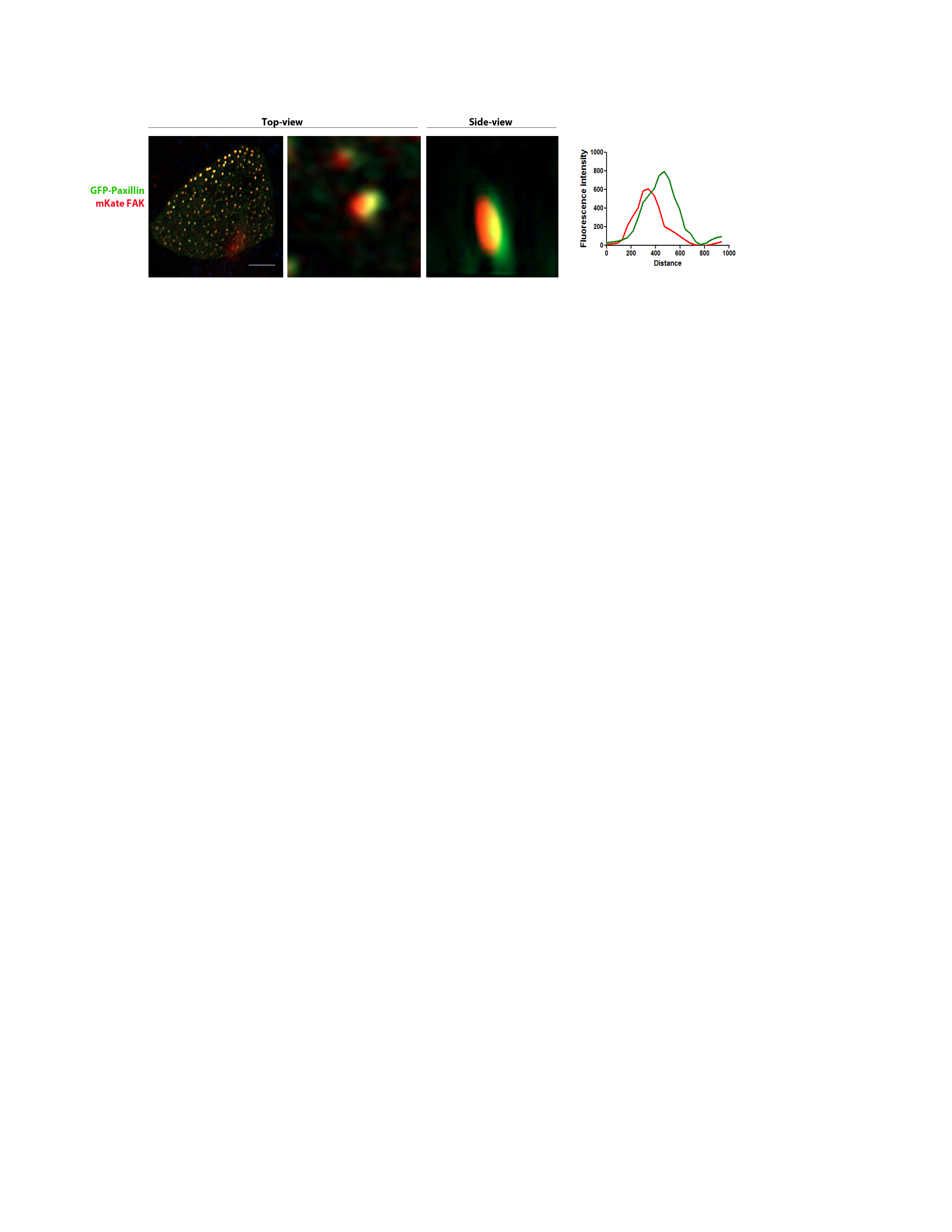
